# SIX4 Controls STING Expression Enhancing anti-PD-1 Efficacy

**DOI:** 10.1101/2023.05.26.542458

**Authors:** Beiyuan Liang, Evan H. Zhang, Zhen Ye, Hayden Storts, Xinru Zheng, Olivia Zaleski, Wei Jin, Xuanxuan Xing, Jing J. Wang

## Abstract

The cGAS/STING cytosolic DNA sensing pathway plays a central role in anti-tumor immunity. Expression of STING is tightly regulated and commonly reduced or defective in many types of cancer. We have identified SIX4 as a significant regulator of STING expression in colon cancer cells. We showed that knockout of SIX4 decreased STING expression at the mRNA and protein levels while ectopic expression of SIX4 increased STING expression. Depletion of SIX4 led to attenuated STING activation and downstream signaling. Re-expression of SIX4 or ectopic expression of STING in SIX4 knockout cells reversed the effect. Ectopic expression of SIX4 enhanced DMXAA and cGAMP-induced STING activation and downstream signaling. Importantly, decrease of SIX4 expression substantially decreased tumor infiltration of CD8^+^ T-cells and reduced the efficacy of PD-1 antibodies to diminish tumor growth in immune competent mice *in vivo*. Finally, analysis of TCGA colon cancer dataset indicated that tumors with high SIX4 expression were significantly enriched in the Inflammatory Response pathway. SIX4 expression also correlated with expression of multiple IFN-stimulated genes, inflammatory cytokines and CD8A. Taken together, our results implicate that SIX4 is a principal regulator of STING expression in colon cancer cells, providing an additional mechanism and genetic marker to predict effective immune checkpoint blockade therapy responses.

## Introduction

Cyclic GMP-AMP (cGAMP) synthase (cGAS) is a cytosolic DNA sensor that when activated generates cGAMP. cGAMP is a second messenger that triggers the stimulator of interferon genes (STING) adaptor to activate downstream signaling, including the Tank-binding kinase-1 (TBK1)/IRF3 and the IkB kinase (IKK) of the NF-kB signaling. Activation of IRF3 and NF-kB induce the expression of type I interferons and inflammatory cytokines and chemokines to initiate innate and adaptive immune responses ^1^. Numerous studies have demonstrated that DNA damage, genomic instability and cellular stress activate cGAS/STING signaling ^2, 3^. Moreover, tumor cell-intrinsic cGAS/STING activation has been linked to efficient cancer treatment including radiation, chemotherapies and anti-PD-1/PD-L1 therapies ^4, 5^, possibly attributed to increased infiltration of T-cells and/or enhanced tumor cell recognition by T-cells ^5, 6^. Interestingly, the cGAS/STING pathway is frequently suppressed in a variety of cancers ^5, 7^. The identification of strategies that restore or reactivate tumor cell-intrinsic cGAS/STING signaling may be useful to enhance cancer treatment efficacy.

Because it is an important regulator of innate immunity, STING expression is tightly controlled. Previous work has demonstrated that STING is regulated at the transcriptional, post-transcriptional, and post-translational levels ^6-11^. For example, GATA1 and Sp3 have been shown to bind to the STING promoter and modify its transcription in NIH3T3 cells ^8^. Hypermethylation of the STING promoter is responsible for silenced STING expression in melanoma and colon cancers ^6, 7^. In addition, microRNA-576-3p has been identified as a negative regulator of STING expression that enables vesicular stomatitis virus (VSV) replication ^9^. Moreover, STING protein stability is regulated by ubiquitination, where USP18/USP20 increases STING stability by reducing its ubiquitination ^10^. STING activation has also been shown to be regulated by phosphorylation ^11^. Despite these studies, the regulation of STING expression *in vivo* remains largely unknown.

Sine oculis homeobox (SIX) 4 is one of the SIX family of homeobox transcription factors ^12^. SIX4 expression is upregulated and has been shown to play an important role in tumorigenesis and metastasis of lung, breast, colorectal cancer ^13-15^. SIX4 regulates transcription of many target genes, such as YAP1 and c-MET in hepatocellular carcinoma and IDH1 in osteosarcoma ^16, 17^. Moreover, SIX4 promotes PI3K/AKT activation in colorectal cancer ^14^ and activates STAT3 in breast cancer ^13^.

Here we show that SIX4 is a major regulator of STING expression in colon cancer cells. Knockout of SIX4 decreased STING expression at the mRNA and protein levels, whereas ectopic expression of SIX4 increased STING expression. Reduced SIX4 expression resulted in attenuated activation of STING signaling by STING agonists. Re-expression of SIX4 or ectopic expression of STING in SIX4 knockout cells reversed the effect. As expected, ectopic expression of SIX4 enhanced STING activation. Depletion of SIX4 significantly reduced efficacy of tumor clearance mediated by an anti-PD-1 antibody in syngeneic immune competent mice *in vivo*. The reduction of SIX4-dependent tumor clearance was associated with decreased tumor infiltration of CD8^+^ T-cells. Analysis of TCGA colon cancer dataset is consistent with a role for SIX4 in controlling STING-dependent tumor clearance. Taken together, our studies demonstrate that SIX4 increases STING expression and activation in colon cancer cells, providing an additional mechanism and genetic marker to predict effective response of colon cancer patients to immune checkpoint blockade therapies.

## Materials and Methods

### Colon cancer cells

Mouse colon cancer cell lines, MC38 and CT26, and human colon carcinoma cell line, HT29, were purchased from ATCC. Human colon carcinoma cell line, TENN, was established in Dr. Brattain’s lab in 1981 ^18^. All cell lines were authenticated by STR analyses at Ohio State University Genomics Shared Resources. STR profiles were cross-checked with the ATCC database. MC38, CT26 and HT29 cell lines displayed ≥ 80% match, which is considered valid ^19^. STR profile of TENN cell line has never been reported before. It was confirmed to be of human origin and contain no interspecies contamination.

Cells were maintained at 37°C in a humidified incubator with 5% CO2. MC38 and CT26 cells were cultured in DMEM supplemented with 10% FBS while HT29 and TENN cells in McCoy’s 5A medium (Cytiva) with 10% FBS.

### Antibodies and reagents

Anti-SIX4, anti-ISG15 and anti-Actin were purchased from Santa Cruz Biotechnology. Anti-STING, anti-phospho-STING, anti-TBK1, anti-phospho-TBK1, anti-STAT1 and anti-phospho-STAT1 were obtained from Cell Signaling Technology. Mouse STING activator DMAXX and cGAMP were purchased from InvivoGen.

### Real time Q-PCR

Q-PCR analysis was performed using PowerUp™ SYBR™ Green Master Mix (Thermo Fisher). Primer sequences for mouse IFNβ, CXCL10 and STING are ATGAGTGGTGGTTGCAGGC-F, TGACCTTTCAAATGCAGTAGATTCA-R; AGTAACTGCCGAAGCAAGAA-F, GCACCTCCACATAGCTTACA-R and GGAACACCGGTCTAGGAAGC-F, TGGATCCTTTGCCACCCAAA-R respectively. Primer sequences for human IFNβ, CXCL10 and STING are ATGACCAACAAGTGTCTCCTCC-F, GGAATCCAAGCAAGTTGTAGCTC-R; AGCAGAGGAACCTCCAGTCT-F, ATGCAGGTACAGCGTACAGT-R and CTTCACTTGGATGCTTGCC-F, CCCGTAGCAGGTTGTTGTAATG-R respectively. Actin was used as an endogenous control.

### CRISPR/CAS9 knockout of SIX4

The Alt-R CRISPR-Cas9 System (Integrated DNA Technologies) was used, in which guide RNA contains crRNA with specific DNA target sequence and tracrRNA labeled with ATTO™ 550 (ATTO-TEC). Two crRNAs targeting SIX4 (ACAACTCCACTCGGAACTTC and CCTCGCACACGCAGGCGACA) were synthesized. Guide RNAs complexed with CAS9 protein were transfected into MC38 cells by electroporation. ATTO™ 550-positive cells were sorted into pools by flow cytometry.

### Plasmids

Mouse SIX4 cDNA (Genomics-online) was cloned into pCDH-CMV-MCS-EF1α-Puro Lentivector (System Biosciences). Human SIX4 cDNA was amplified from genomic DNA by PCR and cloned into pCDH-CMV-MCS-EF1α-Puro vector.

### *In vivo* xenograft model

Mouse experiments were approved by Ohio State University IACUC. Exponentially growing MC38 cells (0.5 × 10^6^) were inoculated subcutaneously into the flank of 6-week-old C57BL/6 mice (Jackson Laboratory). Mice were randomly divided into two groups, with one group treated with 200 μg of control IgG and the other with anti-PD-1 (Ichorbio) by intraperitoneal injection on day 7, 10 and 13. Tumors were measured every other day.

### Immunohistochemistry (IHC) staining

Formalin-fixed paraffin-embedded blocks of tumors were cut into 4-micron thick tissue sections. After antigen retrieval using Antigen Unmasking Solution (Vector Laboratories), tissue slides were blocked with 10% Goat Serum (Vector Laboratories) diluted in 1X PBS, followed by incubation with an anti-CD8 antibody (Cell Signaling) overnight at 4°C. The slides were developed with ImmPACT**®** DAB Substrate Kit (Vector Laboratories) and counterstained with hematoxylin. Five fields/slide were randomly selected for quantification by ImageJ.

### Bioinformatics analysis of clinical data

RNA-seq data for TCGA-COAD was accessed through GDC data portal (https://portal.gdc.cancer.gov/). TCGA-COAD samples (n=471) were classified in SIX4^high^ or SIX4^low^ based on SIX4 expression using z-score. GSEA was performed on GenePattern (https://cloud.genepattern.org/) using MSigDb genesets ^20^. Correlation of SIX4 with other genes was calculated with Spearman’s correlation. Analysis was conducted using R and RStudio.

### Statistical analysis

Experiments were performed a minimum of two times independently and represented by Mean ± SD. Statistics analysis was performed in GraphPad Prism 5. Unpaired Student’s t-test was used to analyze the differences among groups. Statistically significant differences are indicated as follows: \**P*<0.05, \*\**P*<0.01, \*\*\**P*<0.001.

## Results and Discussion

### SIX4 regulates STING expression and enhances the activation of STING/TBK1/IFNβ signaling

To identify transcriptional regulators of STING, we employed JASPER to screen for STING regulators. These studies suggested that SIX4 might regulate STING expression. We used CRISPR/Cas9 and two distinct guide RNAs (gRNAs) to knockout SIX4 expression in MC38 cells. Deletion of SIX4 significantly reduced STING expression at both mRNA and protein levels, and re-expression of SIX4 rescued STING expression (Fig. 1A). Complementary studies showed that ectopic expression of SIX4 increased STING expression in CT26, HT29 and TENN cells (Fig. 1B).

**Figure 1.**
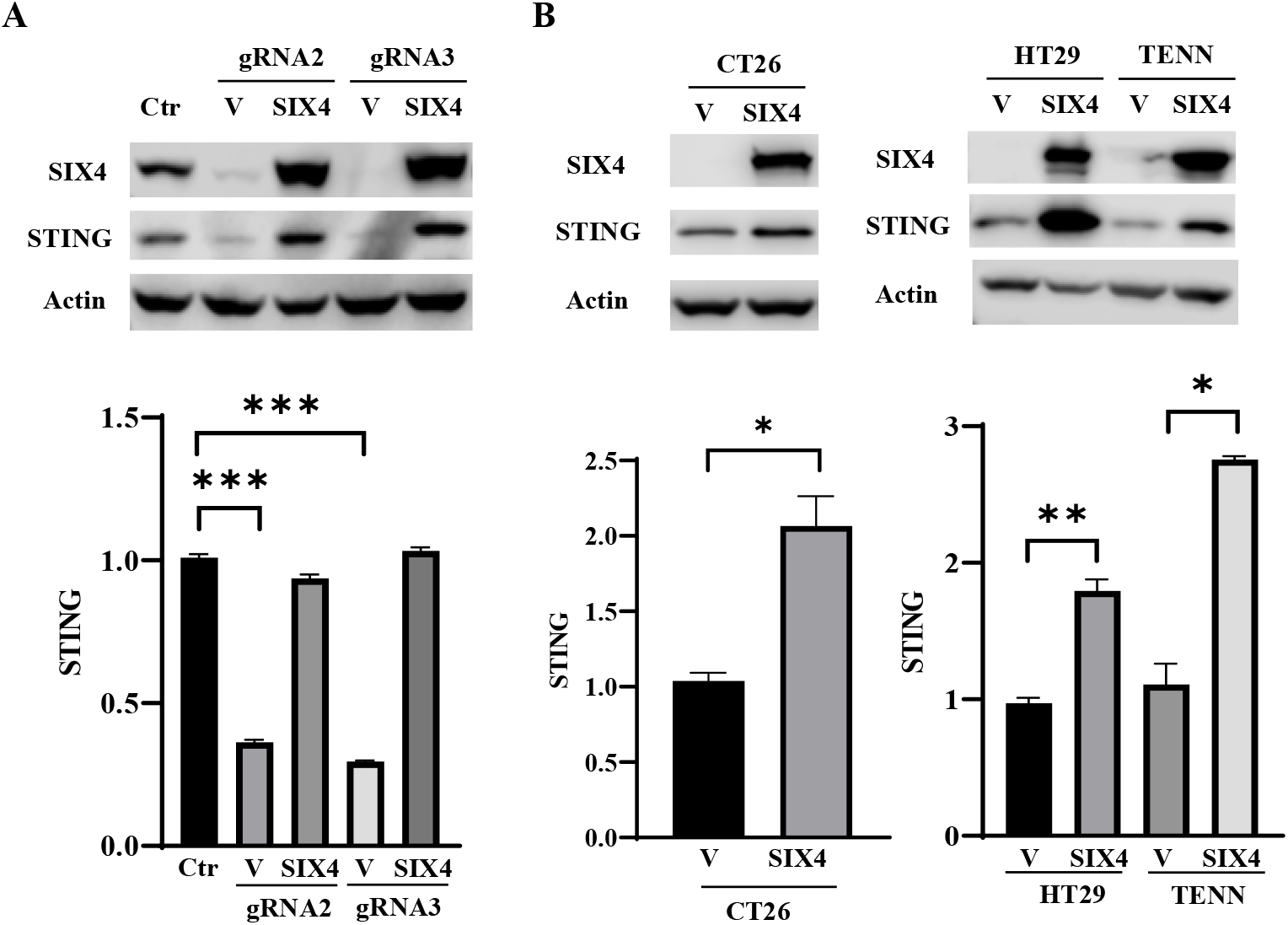
SIX4 regulates expression of STING mRNA and protein. A, SIX4 expression was knocked out in MC38 cells by two gRNAs. SIX4 was then re-expressed in SIX4 knockout cells. Depletion of SIX4 significantly reduced STING expression and re-expression of SIX4 rescued STING expression at mRNA and protein levels determined by western blot (upper panel) and Q-PCR (lower panel) analysis respectively. B, SIX4 was ectopically expressed in CT26, HT29 and TENN cells, which led to increased expression of STING mRNA and protein. Results are shown as mean ± SD. *P < 0.05, **P < 0.01, ***P < 0.001.

A major consequence of STING activation is the production of type I interferons and inflammatory cytokines ^1^. Treatment of MC38 cells with DMXAA, a specific STING agonist, resulted in significant upregulation of STING and TBK1 phosphorylation as well as elevated expression of IFNβ, CXCL10 and ISG15, an IFN-regulated gene (Fig. 2A). Remarkably, total STING protein level was significantly reduced by DMXAA treatment (Fig. 2A). These observations are consistent with the hypothesis that acute activation of STING leads to reduced STING expression to counteract sustained STING activation. Depletion of SIX4 attenuated DMXAA-mediated STING activation, while re-expression of SIX4 or STING reinstated DMXAA effect (Fig. 2A). Conversely, ectopic expression of SIX4 in CT26 cells further enhanced DMXAA-mediated activation of STING signaling and increased expression of IFNβ, CXCL10 and ISG15 (Fig. 2B).

**Figure 2.**
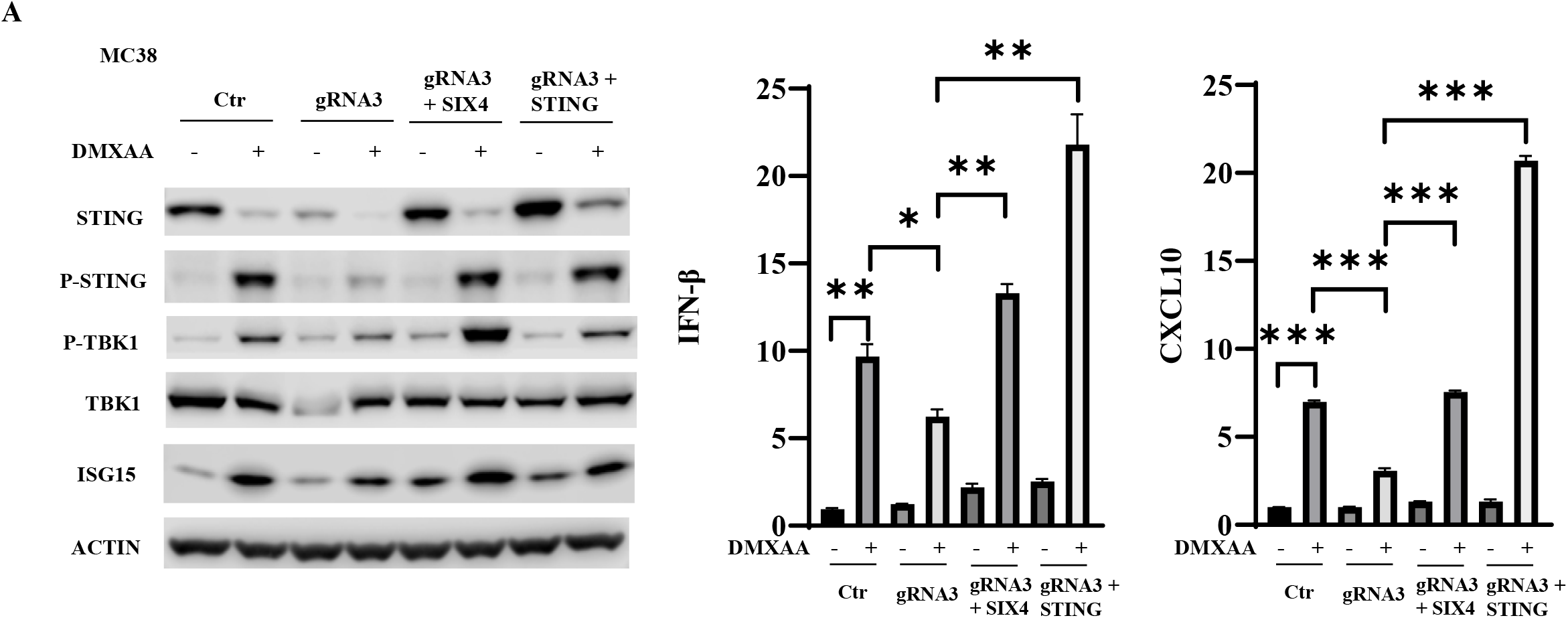
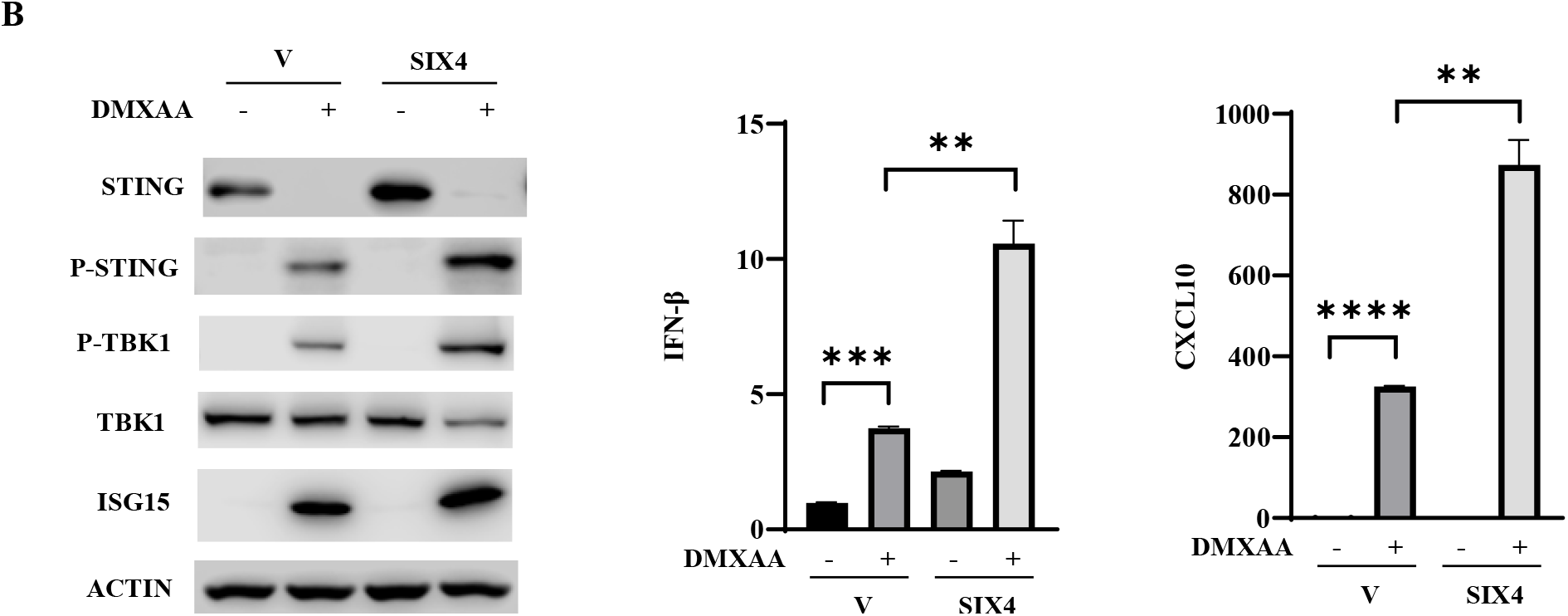
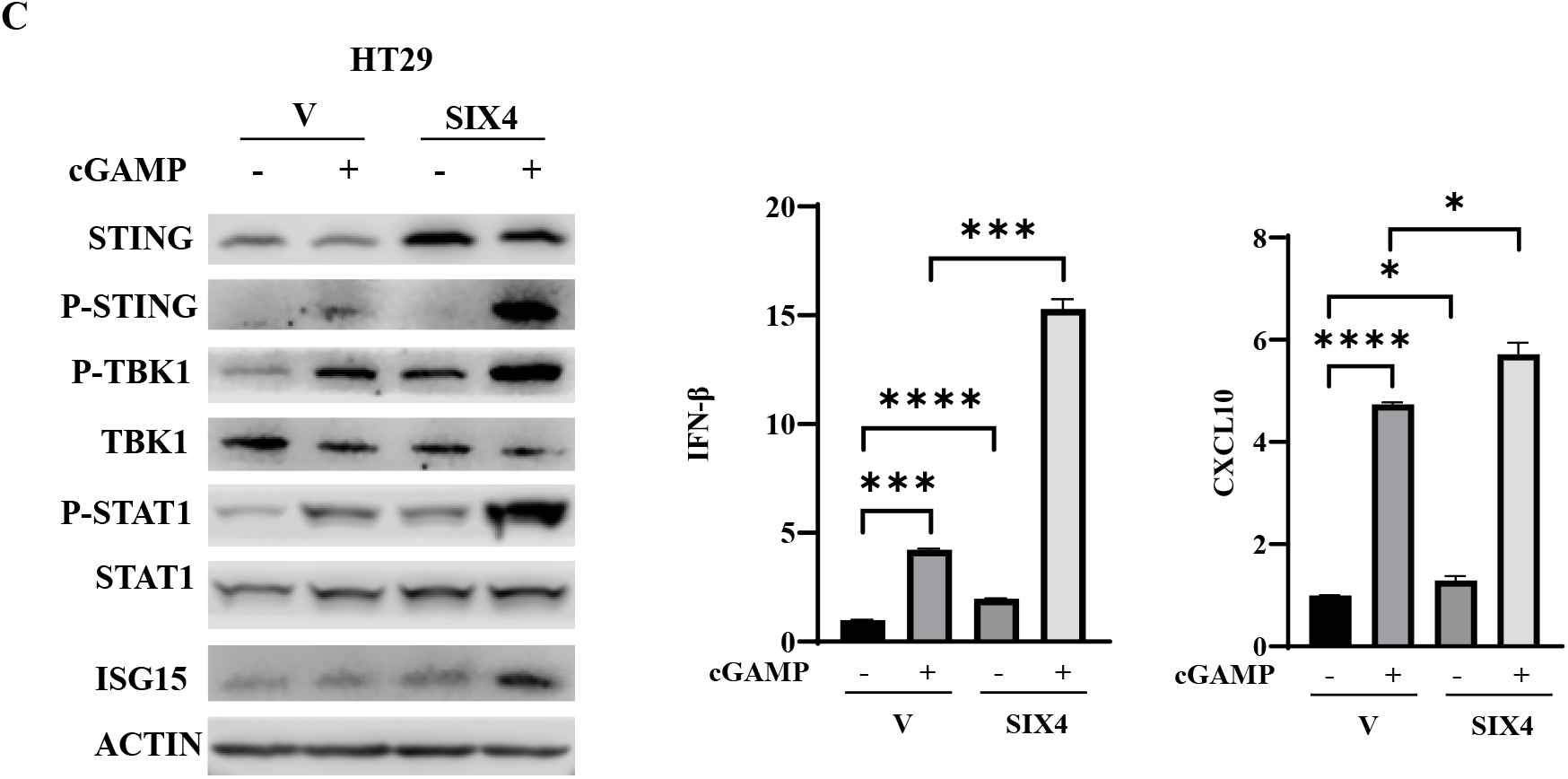
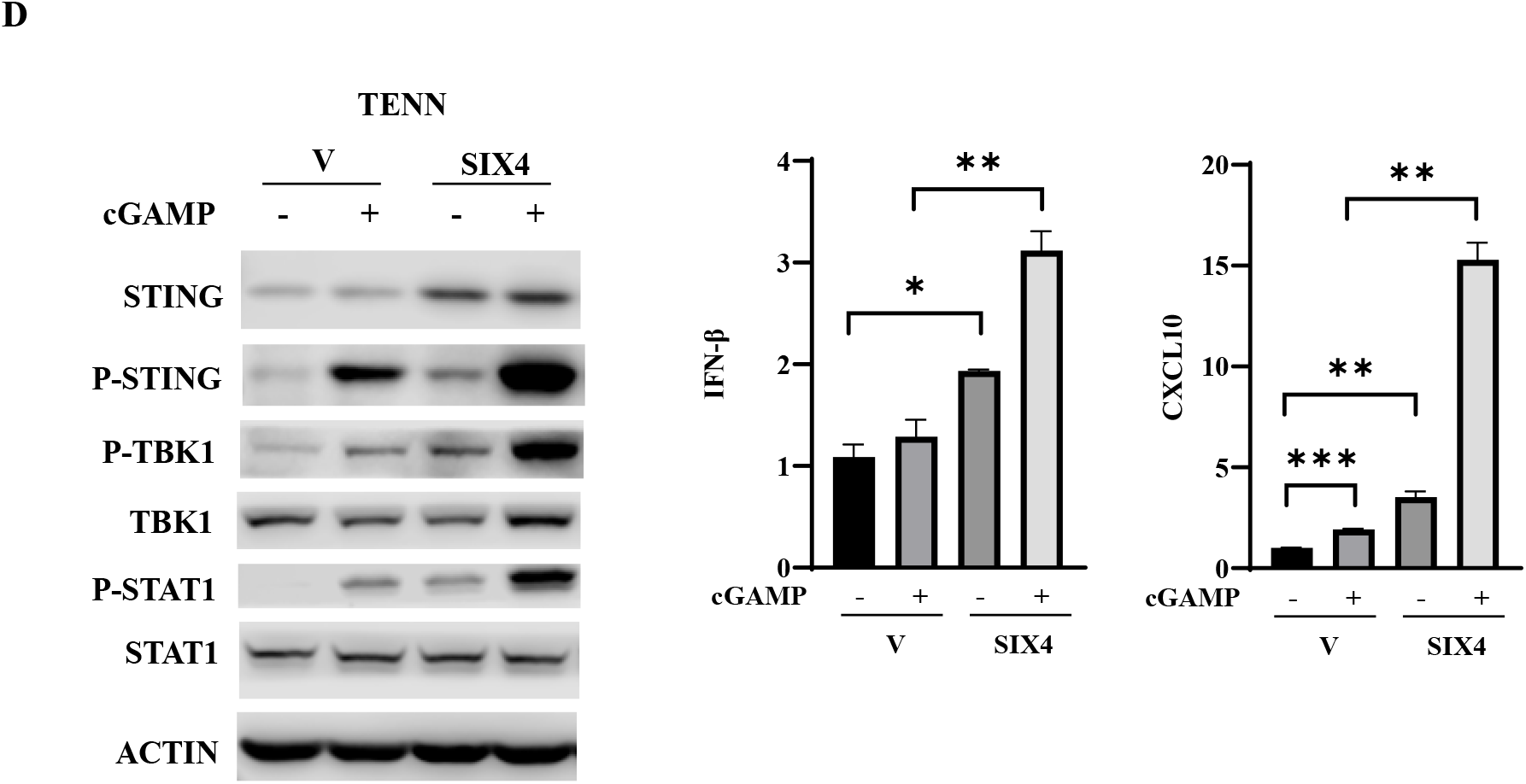
SIX4 modulates STING activation in colon cancer cells. A, MC38 SIX4 knockout cells and those with SIX4- or STING-re-expressing cells were treated with 10 μM DMXAA for 6 hrs. Western blot analysis was performed to examine expression and phosphorylation of STING and TBK1 and ISG15 expression (left panel). IFNβ and CXCL10 mRNA expression was determined by Q-PCR analysis (right panels). B, CT26 control and SIX-overexpressing cells were treated with 10 μM DMXAA for 6 hrs. Western blot analysis was performed to examine expression and phosphorylation of STING and TBK1 and ISG15 expression (left panel). IFNβ and CXCL10 mRNA expression was determined by Q-PCR analysis (right panels). C&D,3 the control and SIX4-overexpressing HT29 and TENN cells were transfected with 10 μg cGAMP. Cells were harvested for protein and RNA isolation. Western blot analysis was performed to examine expression and phosphorylation of STING, TBK1 and STAT1 and ISG15 expression (left panel). IFNβ and CXCL10 mRNA expression was determined by Q-PCR analysis (right panels). Results are shown as mean ± SD. *P < 0.05, **P < 0.01, ***P < 0.001.

To determine whether SIX4 regulates STING/TBK1/IFNβ signaling in human colon cancer cells, HT29 and TENN cells were transfected with cGAMP to activate STING in the presence or absence of SIX4 overexpression. Increased expression of SIX4 enhanced well-known cGAMP-mediated increases in STING, TBK1 and STAT1 phosphorylation as well as upregulation of IFNβ, CXCL10 and ISG15 expression (Fig. 2C and 2D). Taken together, these results indicate that SIX4 activates STING/TBK1/IFNβ signaling through upregulation of STING expression.

### Reduction of SIX4 expression reduces anti-PD-1 mediated suppression of tumor growth in vivo

Previous work has suggested SIX4 enhances cell proliferation in tumors ^12^. To establish whether SIX4 expression affects cell proliferation, we performed colony formation analysis in control, SIX4 knockout and SIX4-overexpressing MC38 cells. The results revealed that depletion or overexpression of SIX4 had little effect on the clonogenicity of MC38 cells in culture (Fig. 3A and 3B).

**Figure 3.**
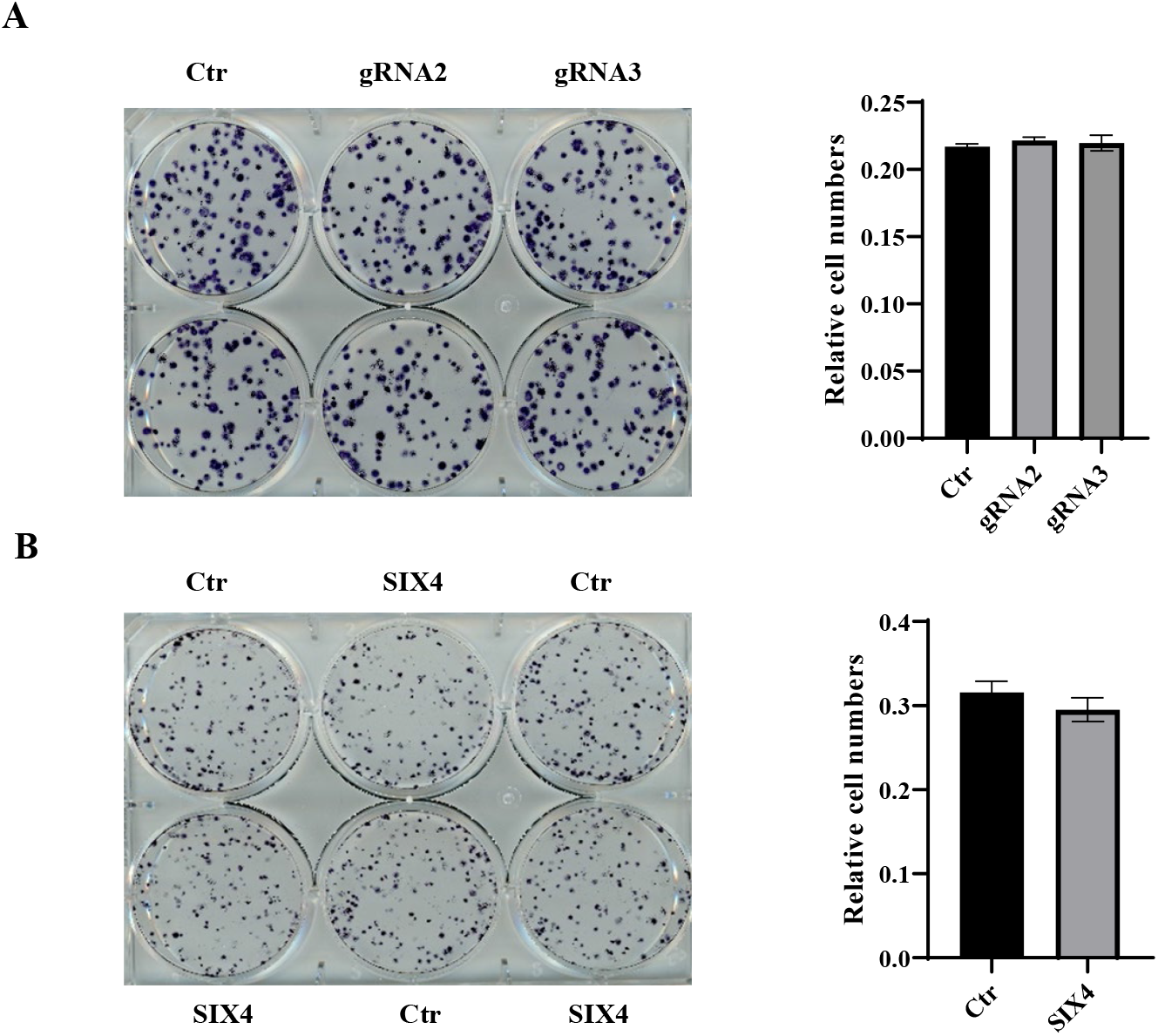
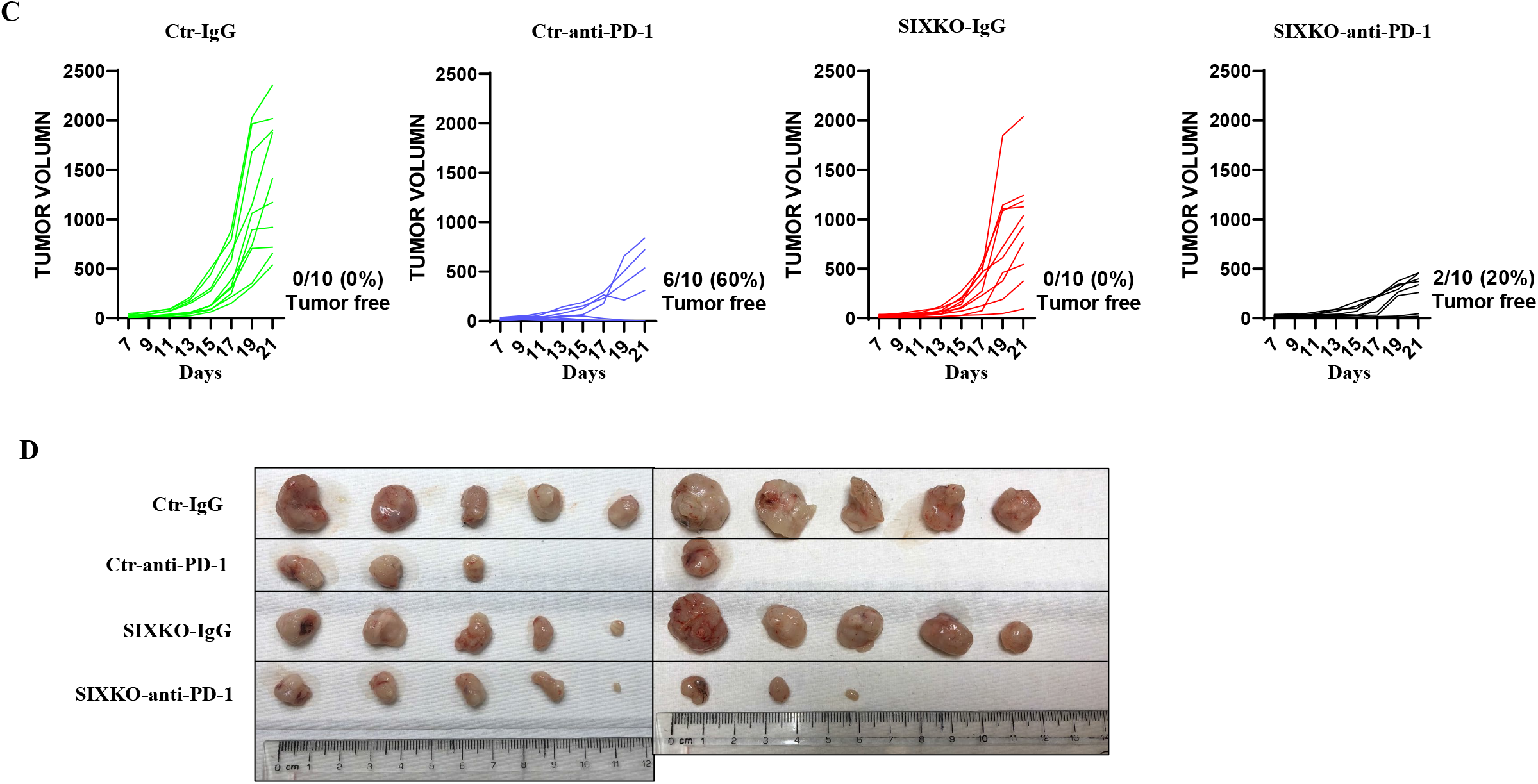
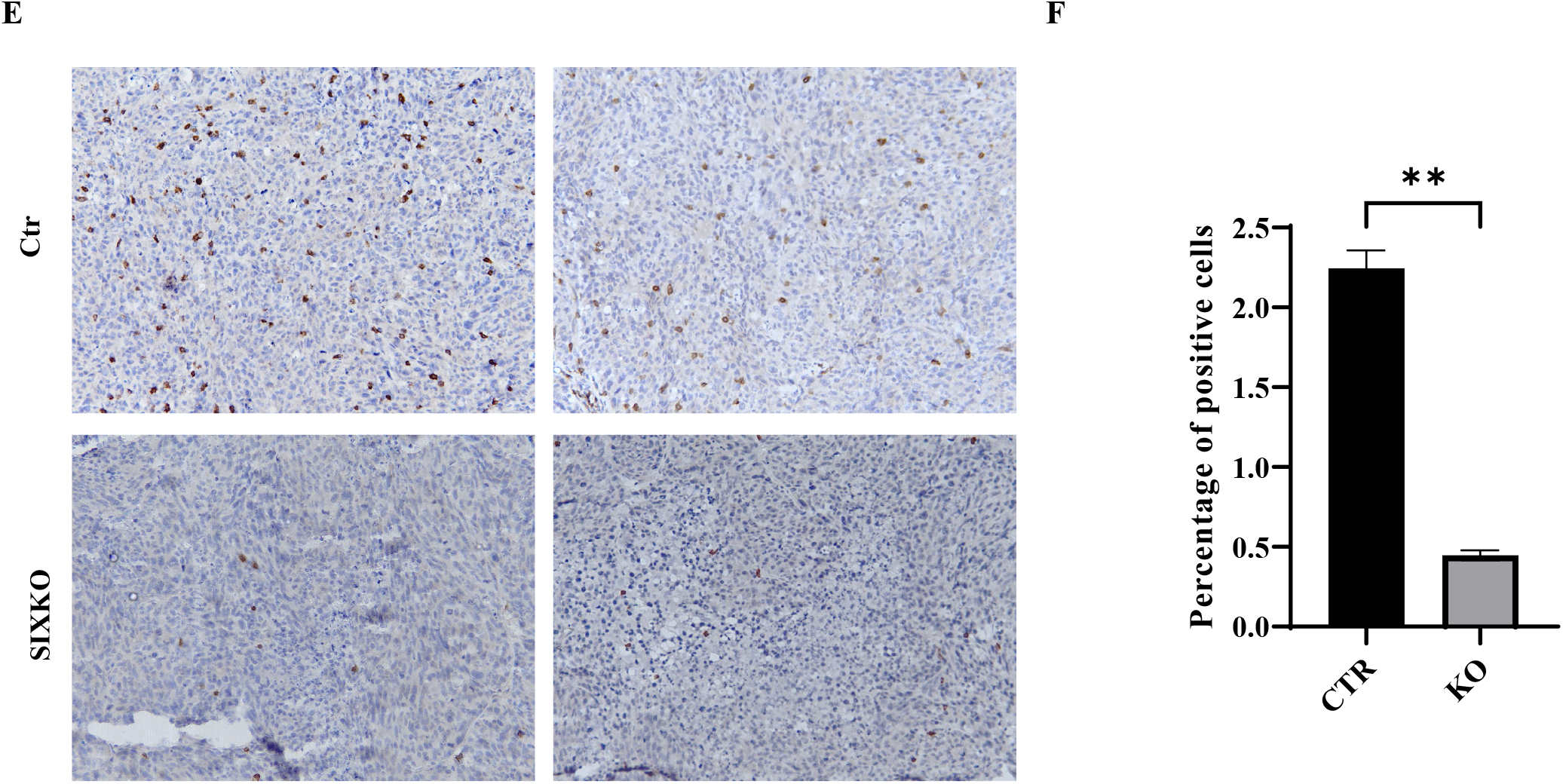
Knockout of SIX4 reduces the efficacy of anti-PD-1 tumor suppression effect *in vivo*. A, Colony formation assays were performed in MC38 control and SIX4 knockout cells (gRNA2 and gRNA3) with representative images (left panel) and quantification (right panel). B, Colony formation assays were performed in MC38 control and SIX4 overexpressing cells with representative images (left panel) and quantification (right panel). C, Xenograft tumor growth curves of MC38 control and SIX4 knockout (KO) cells treated with a control IgG or an anti-PD-1 antibody are shown. N = 10. D, Images of tumors at the endpoint of experiments are shown. E, Representative 200x images of IHC staining of CD8 in control and SIX4 knockout tumors. F, Quantification of percentage of CD8-positive cells. Results are shown as mean ± SD. **P < 0.01.

Given the importance of STING activation in the efficacy of anti-PD-1/PD-L1 therapies ^4^, we next determined the effect of SIX4 expression on the efficiency of an anti-PD-1 antibody-mediated inhibition of tumor growth in immune competent mice. Control and SIX4 knockout MC38 cells were injected subcutaneously into *wild type* C57BL/6 mice that were then randomly divided into two groups and treated with a normal IgG or an anti-PD-1 antibody respectively. Although depletion of SIX4 did not have a significant effect on tumor growth, it substantially reduced the response to anti-PD-1 treatment. Treatment with normal IgG in both control and SIX4 knockout tumors resulted in 100% tumor growth. Notably, treatment with the anti-PD-1 antibody led to complete disappearance of tumors in 6 out 10 mice bearing control cells whereas it only cleared tumors in 2 out of 10 mice bearing SIX4 knockout cells (Fig. 3C and 3D).

One of the mechanisms by which tumor cell-intrinsic STING activation contributes to effective anti-PD-1/PD-L1 therapies is by increasing intra-tumoral T-cells ^5^. IHC staining of CD8 in tumor sections showed a marked reduction of CD8-positive T-cells in the tumors of SIX4 knockout cells as compared to control tumors (Fig. 3E). Quantification indicated that percentage of CD8^+^ T cells in SIX4 knockout tumors was reduce to 22% of that in control tumors (Fig. 3F). Taken together, these results indicate that depletion of SIX4 reduced CD8^+^ T-cell infiltration and attenuated tumor response to anti-PD-1 treatment.

### SIX4 expression is associated with inflammatory response in colon cancer patient specimens

We examined the relationship between SIX4 expression and STING-dependent inflammatory activities in human colon cancer by mining RNA-seq data from TCGA colon cancer dataset (TCGA-COAD). Patients were stratified into SIX4 high and low groups. Differential gene expression was determined between two groups and GSEA was performed. SIX4 high group was significantly enriched in the Inflammatory Response pathway (Fig. 4A). In addition, SIX4 expression was highly correlated with expression of multiple ISGs (Fig. 4B), inflammatory cytokines (Fig. 4C) and CD8A that is predominantly expressed on cytotoxic T-cells (Fig. 4D). These results suggest that high SIX4 expression in colon cancer patients likely results in enhanced STING activation, inflammatory response, and increased infiltration of CD8^+^ lymphocytes as compared to low SIX4 expression. These results appear largely identical to our *in vitro* and *in vivo* results using colon cancer cell lines.

**Figure 4.**
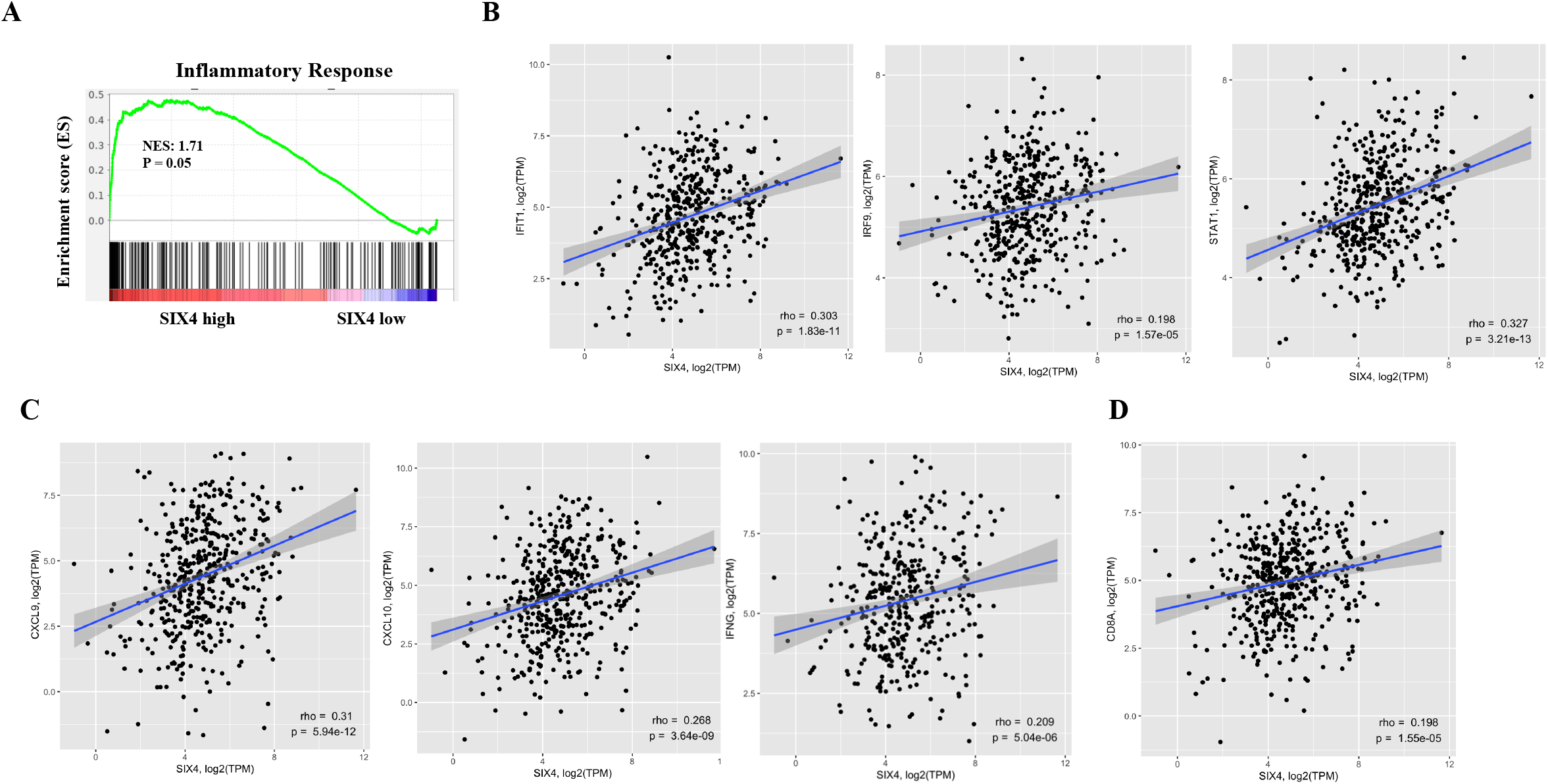
SIX4 expression is positively associated with inflammatory response in colon cancer patient specimens. A, GSEA plots of enrichment in the Inflammatory Response signatures from MsigDB in colon cancer patient samples from TCGA stratified by SIX4 expression. B, Correlation between SIX4 and IFN marker gene expression in colon cancer patient samples from TCGA. C, Correlation between SIX4 and inflammatory cytokine/chemokine expression in colon cancer patient samples from TCGA. D, Correlation between SIX4 and CD8A expression in colon cancer patient samples from TCGA.

### Tumor suppression function of SIX4

Previous studies have suggested that SIX4 function is tumor-promoting ^13-17^. In contrast, our studies demonstrate that SIX4 enhances STING/IFNβ innate immune signaling by upregulating STING expression. Tumor cell intrinsic STING activation plays an important role in anti-tumor immunity by increasing tumor infiltrating T cells, reducing tumor-associated myeloid cells and/or enhancing tumor antigenicity and T cell recognition ^5, 6^. This tumor suppressing activity of SIX4 adds an additional dimension to the SIX4 function.

SIX4 expression has been shown to be upregulated in colorectal cancer ^14^. High SIX4 expression is likely to increase STING expression based on our studies. However, it appears that STING promoter is hypermethylated in some colorectal tumors, leading to silencing of STING expression ^7^. In STING-silenced tumors, SIX4 would not be able to upregulate STING expression, in which case treatment with demethylating agents such as azacytidine or decitabine may restore upregulation of STING expression by SIX4. These observations strongly suggest that tumor cells may evolve multiple mechanisms to escape SIX4’s tumor suppression function ultimately facilitating its tumor promoting activity. Our analysis of colon cancer patient data indicated that tumors with high SIX4 expression were significantly enriched in the Inflammatory Response pathway and that SIX4 expression was positively associated with CD8A expression. It seems likely that colon cancer patients with high SIX4 expression may have immunogenic tumors with infiltration of CD8^+^ T-cells. These patients are likely to respond well to the anti-PD-1/PD-L1 treatment. Furthermore, for those patients with methylated STING promoter, the combination of anti-PD-1/PD-L1 with demethylating agents (e.g., azacytidine or decitabine) would likely improve the treatment efficacy.

In summary, we have identified SIX4 as a novel regulator of STING expression, which has a tumor suppression function by increasing anti-tumor immunity. Knockout of SIX4 attenuated STING activator-mediated activation of STING/IFNβ signaling cascade and reduced efficacy of an anti-PD-1 antibody to eliminate tumor growth in immune competent mice. Data analysis of human colon cancer patients support the hypothesis that SIX4 increases STING/type I IFN signaling and enhances inflammatory responses, which facilitates T cell infiltration and anti-tumor immunity. SIX4 could regulate STING expression directly by binding to its promoter or indirectly through its target genes. Nevertheless, SIX4 regulation of STING expression is significant in colon cancer. Therefore, our studies add an important layer to SIX4 function in regulating colon cancer progression and identify it as a potential target for enhancing immunotherapy.

## Acknowledgements

The results shown here are in whole or part based upon data generated by the TCGA Research Network: https://www.cancer.gov/tcga. The work is supported by NIH grants R01CA215389 and R01CA208063 to JW, by the Pelotonia Institute of Immuno-Oncology (PIIO), and by The Ohio State University Comprehensive Cancer Center and the National Institutes of Health under grant number P30 CA016058. We thank Dr. Richard Fishel for discussion and editing of the manuscript and Dr. Wayne Miles for discussion and help. We also thank the Genomics and Flow Cytometry Shared Resources at The Ohio State University Comprehensive Cancer Center.

